# Unstable Angina is a syndrome correlated to Th17 inflammatory disorder

**DOI:** 10.1101/396267

**Author notes:** Correspondence to: Department of Clinical Pathology Far Eastern Memorial Hospital No. 21, Nan-Ya South Road section2, Ban-Ciao District, New Taipei City, Taiwan.

## Abstract

**Purpose:** Unstable angina is common clinical manifestation of atherosclerosis. However, the detailed pathogenesis of unstable angina is still not known. Here, I propose that unstable angina is a TH17 dominant inflammatory disorder.

**Methods:** Microarray dataset from unstable angina patients from Gene Expression Omnibus publicly available website is used for further analysis compared to healthy control.

**Results:** I find out that TH17 related cytokine, cytokine receptor, chemokines, complement, immune-related transcription factors, anti-bacterial genes, Toll-like receptors, and heat shock proteins are all up-regulated in peripheral leukocytes of unstable angina. In addition, H^+^-ATPase, glycolytic genes, platelet and RBC related genes are also up-regulated in peripheral leukocytes of during unstable angina. Pathway analysis also supports that TH17 immunological pathway is over-represented in the unstable angina dataset.

**Conclusions:** This finding implies that atherosclerosis is correlated to TH17 inflammatory disease. If we know the etiology of unstable angina as well as atherosclerosis better, we can have better methods to control and prevent this detrimental illness.

## Introduction

Atherosclerosis is a very popular disease, especially in developed countries. It is the top death-causing disease in industrial countries. Coronary artery disease (CAD) is the most important clinical symptom of atherosclerosis. Typical angina and unstable angina are common manifestations of CAD. Atherosclerosis is thought to be an chronic inflammatory disorder. However, the detailed pathogenesis of angina is not known. Here, I use microarray analysis of peripheral leukocytes to describe unstable angina is related to a TH17 inflammatory disorder.

Although atherosclerosis is long thought as an inflammatory disorder, its detailed mediating factors are still unclear. TH17 pathway is a recently identified immunological pathway. It is considered to be a host immunological pathway against extracellular bacteria. Th17 immune response is driven by TGF beta and IL6. Th17 effector cells are neutrophils, IL-17 secreting CD4 T cells, and IgM and IgG secreting B cells. TNF alpha, IL1, IL8, and complements also play important roles in Th17 immunity. Th17 over-activation has been proposed in the pathogenesis of variety of autoimmune diseases. Type 3 immune-complex mediated hypersensitivity is closely related to Th17 immune response. CRP, an important TH17 anti-bacterial player, is usually elevated in atherosclerosis patients’ serum. In this study, I find out that TH17 immunological pathway plays an important role in the pathogenesis of unstable angina which is caused by atherosclerosis.

## Methods

### Study samples

I use microarray dataset from Gene Omnibus website (GEO). The dataset serial number is GSE. This dataset includes peripheral leukocytes from patients of acute myocardial infarction and unstable angina. Whole blood samples were collected in 36 myocardial infarction patients and 16 unstable angina patients on 7 days and 30 days post ACS attacks. Here, I compare the RNA expression profile of leukocytes from unstable angina patients to that from health normal control. This dataset is GSE29111 from Dr. Milo and Rothman’s study group. The healthy normal controls are from dataset of a Huntington’s disease blood biomarker study population. The dataset is GSE8762 from Dr. Runne H.’s study group. This second study was published in PNAS,2007, 104(36):14424.

### RMA normalization & significance analysis

Affymetrix HG-U133A 2.0 genechip was used in both samples. RMA express software(UC Berkeley, Board Institute) is used to do normalization and to rule out the outliners of the above dataset. I removed samples which exceed 99% in RLE-NUSE T2 plot.

By the exclusion criteria above, I removed GSM721017 from unstable angina samples and GSM217774 from healthy normal controls. (Figure 1 & Figure 2) Then, Genespring XI software was done to analysis the significant expressed genes between ARDS and healthy control leukocytes. P value cut-off point is less than 0.05. Fold change cut-off point is >2.5 fold change (either up-regulated or down-regulated). Benjamini-hochberg corrected false discovery rate was used during the analysis. Totally, a genelist of 3388 genes was generated from the HGU133A2.0 chip with 18400 transcripts including 14500 well-characterized human genes.

**Figure 1.**
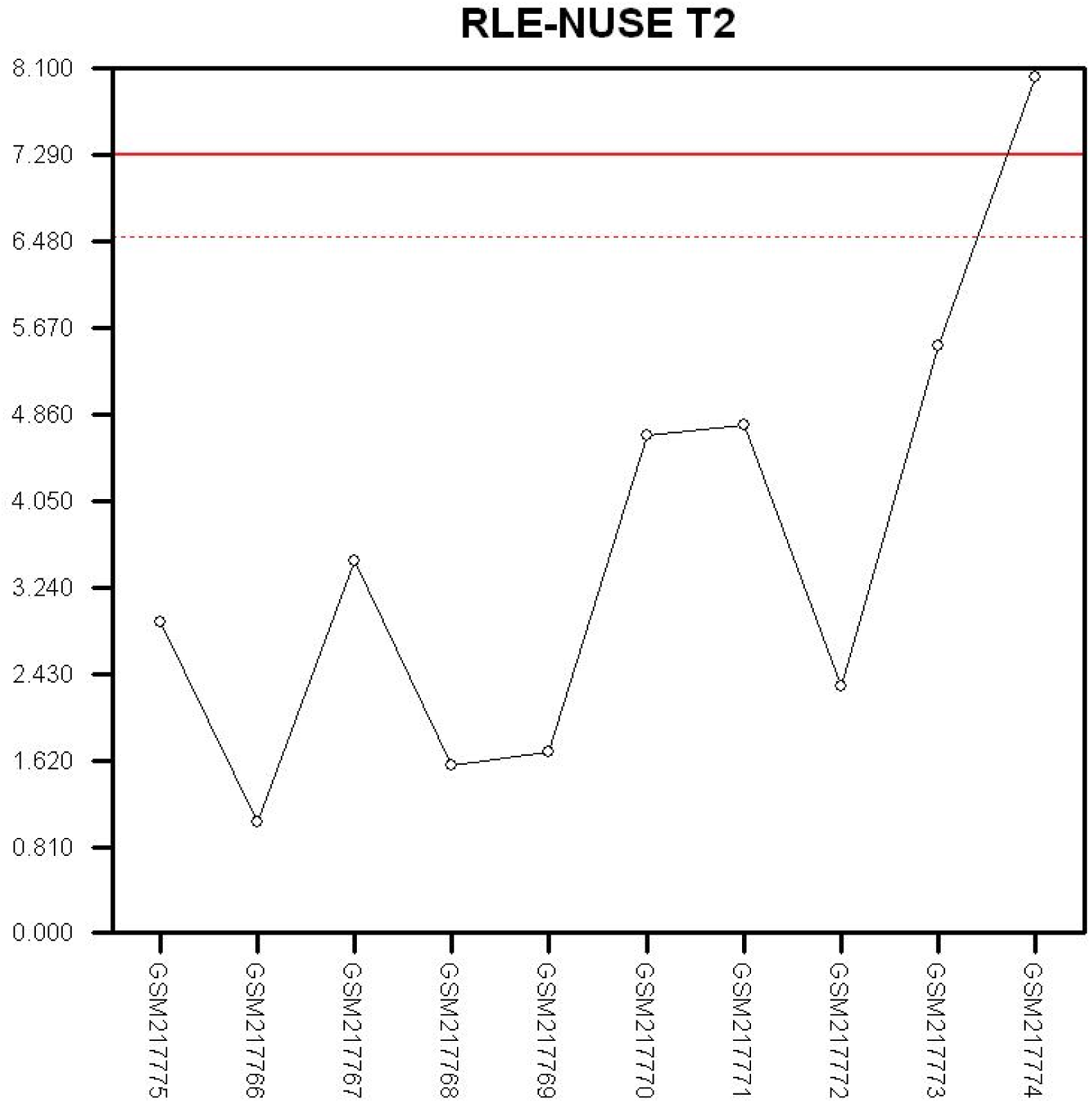
RMA express T2 plot for selecting samples in normal healthy controls. The bold red line is the 99% cutoff line in RLE-NUSE T2 plot.

**Figure 2.**
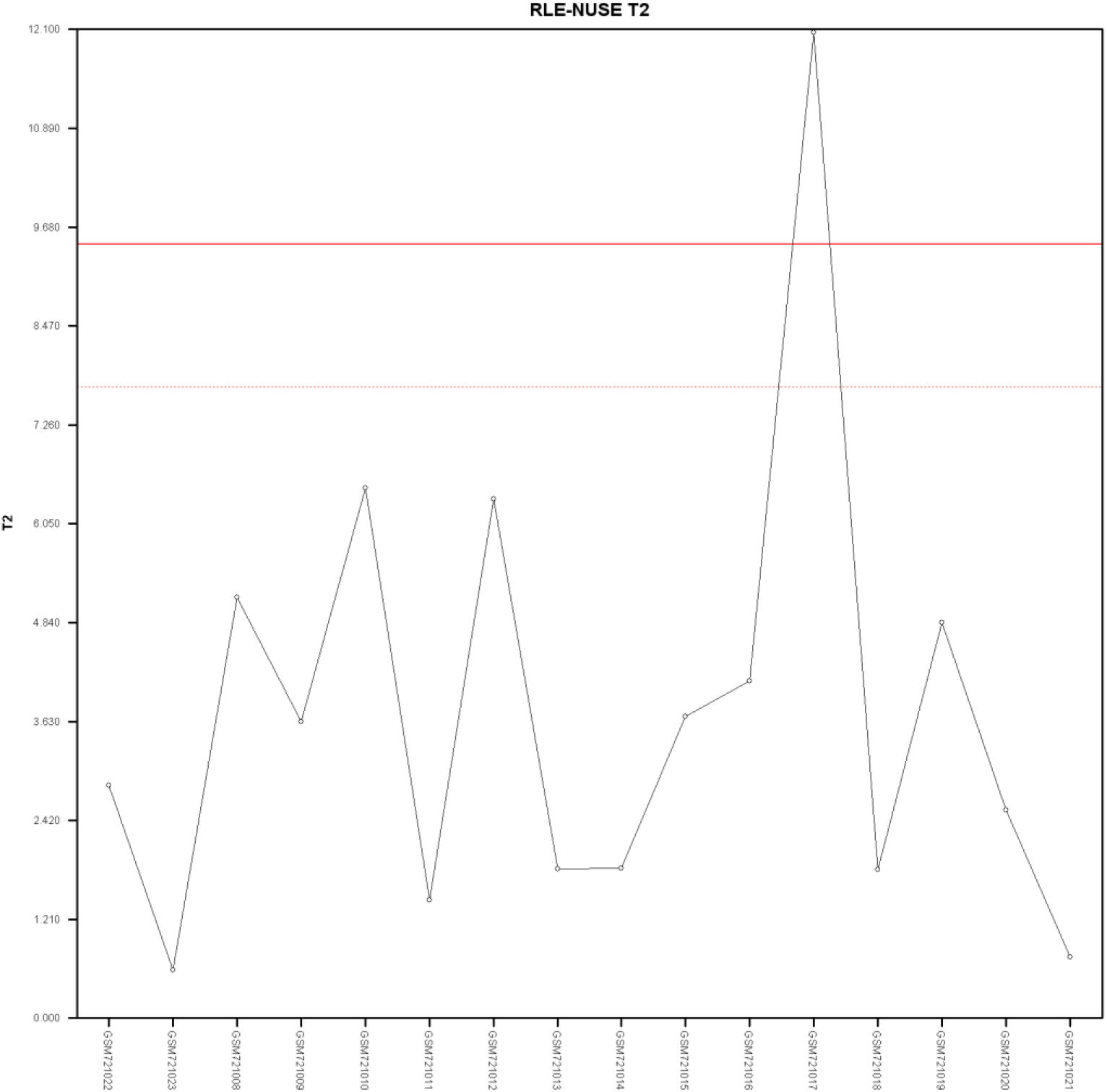
RMA express T2 plot for selecting samples in unstable angina patients. The bold red line is the 99% cutoff line in RLE-NUSE T2 plot.

### Pathway analysis

After genelist of significance analysis was generated. Ingenuity Pathway Analysis were done to identified significantly represented functions or pathways in whole blood cell sample RNAs in unstable angina patients compared to those in normal healthy control. Pathway analysis can help to identify which biological function or pathway is up-regulated or down-regulated in disease pathophysiology.

## Results

### Toll-like receptor up-regulation in unstable angina

During the microarray analysis of peripheral blood leukocytes of unstable angina patients, I find out that Toll-like receptor genes are up-regulated.(Table1) These genes include TLR 1,2,4,8, TIRAP, TTRAP, TANK, and TOLIP. It is worth noting that TLR1 is 7 fold up-regulated, TLR4 is 13 fold up-regulated, TLR2 is 2.7 fold up-regulated, and TANK is 6 fold up-regulated. Since TLR 1,2,4 are related to the activation of antibacterial TH17 immunity, we can say that TH17(anti-extracellular bacteria) immunity is triggered during unstable angina.

**Table 1.**
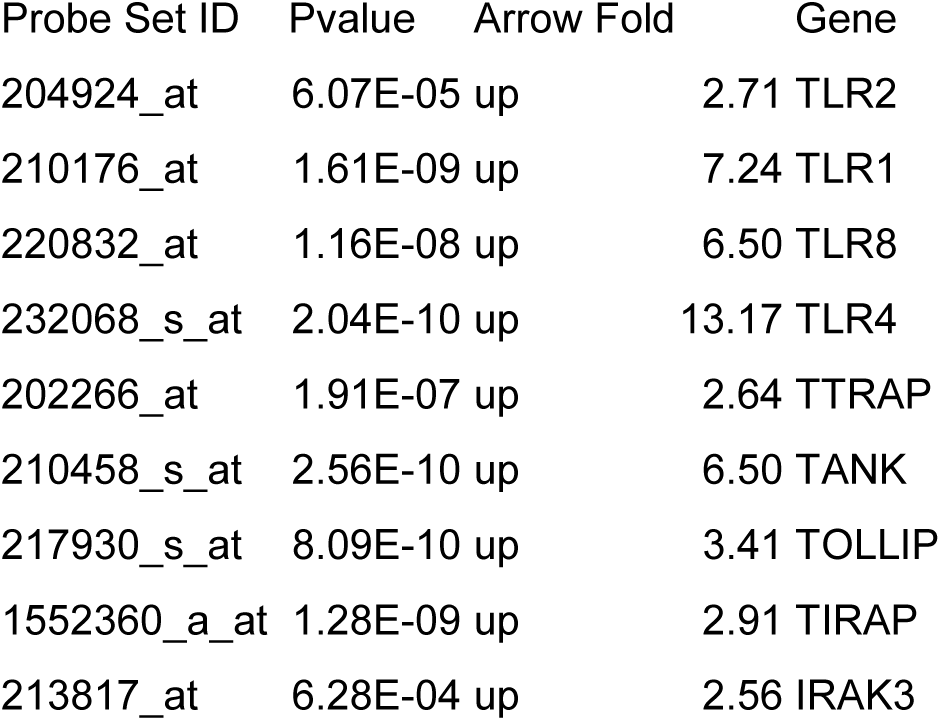
Toll-like-receptor

| Probe Set ID | Pvalue | Arrow | Fold | Gene |
| --- | --- | --- | --- | --- |
| 204924_at | 6.07E-05 | up | 2.71 | TLR2 |
| 210176_at | 1.61E-09 | up | 7.24 | TLR1 |
| 220832_at | 1.16E-08 | up | 6.50 | TLR8 |
| 232068_s_at | 2.04E-10 | up | 13.17 | TLR4 |
| 202266_at | 1.91E-07 | up | 2.64 | TTRAP |
| 210458_s_at | 2.56E-10 | up | 6.50 | TANK |
| 217930_s_at | 8.09E-10 | up | 3.41 | TOLLIP |
| 1552360_a_at | 1.28E-09 | up | 2.91 | TIRAP |
| 213817_at | 6.28E-04 | up | 2.56 | IRAK3 |

### Heat shock protein up-regulation in unstable angina

In the peripheral leukocyte of unstable angina patients, mRNAs of many heat shock proteins are up-regulated. These genes include HSPA6, HSPA1A/A1B, DNAJA4, DNAJB9, DNAJB2, DNAJB4, DNAJC6, and DNAJA2. (Table2) Among them, HSPA1A/HSPA1B is 5.6 fold up-regulated and DNAJC6 is 4.8 fold up-regulated. However, few certain heat shock protein genes are down-regulated including HSP90AA1, DNAJC1, DNAJC2, DNAJC3, and DNAJC7. In addition, these genes are only 2-3 fold down-regulated. HSP70 can bind to TLRs to initiate anti-bacterial immunity. Thus, anti-bacterial TH17 is likely to be activated at unstable angina.

**Table 2.** Heat shock protein

| Probe Set ID | Pvalue | Arrow | Fold | Gene |
| --- | --- | --- | --- | --- |
| 211968_s_at | 5.02E-09 | down | -3.04 | HSP90AA1 |
| 117_at | 2.01E-06 | up | 3.37 | HSPA6 |
| 202581_at | 7.23E-10 | up | 5.62 | HSPA1A/B |
| 1554333_at | 4.79E-05 | up | 4.76 | DNAJA4 |
| 1554462_a_at | 6.27E-08 | up | 2.53 | DNAJB9 |
| 1558080_s_at | 2.64E-06 | down | -3.33 | DNAJC3 |
| 202416_at | 1.77E-09 | down | -2.94 | DNAJC7 |
| 202500_at | 6.74E-09 | up | 3.79 | DNAJB2 |
| 202843_at | 5.11E-09 | up | 2.77 | DNAJB9 |
| 203810_at | 4.20E-08 | up | 2.99 | DNAJB4 |
| 204720_s_at | 2.13E-06 | up | 4.81 | DNAJC6 |
| 209015_s_at | 2.35E-07 | up | 2.84 | DNAJB6 |
| 209157_at | 3.03E-08 | up | 2.65 | DNAJA2 |
| 213097_s_at | 3.59E-10 | down | -2.68 | DNAJC2 |
| 222620_s_at | 5.74E-07 | down | -2.90 | DNAJC1 |

### TH17 related transcription factor up-regulation during unstable angina

Then, we will look at THfh and TH17 related transcription factor regulation during unstable angina.(Table 3) THfh related transcription factors, STAT1, BCL6, and STAT5B are up-regulated during unstable angina. THfh is the initiation of host adaptive immunity. Up-regulated TH17 related transcription factors include SMAD4, SMAD2, SMAD5, SMAD7, RARA, RXRA, STAT5B, FOSL2, and FOXO3. Besides, negative regulator of TH17 immunity, ETS1, is down-regulated in unstable angina. Retinoic acid (for RXRA and RARA) can promote TH17 immunity. SMADs and STAT5B are downstream mediators of TGFb, a key cytokine of TH17 immunity. SOCS3, FOSL2, and FOXO3 can all promote TH17 immunity.

**Table 3.** Transcription factors

| Probe Set ID | Pvalue | Arrow | Fold | Gene |
| --- | --- | --- | --- | --- |
| M97935_MB_at | 5.29E-06 | up | 3.25 | STAT1 |
| 201329_s_at | 2.07E-04 | up | 2.83 | ETS2 |
| 202527_s_at | 2.47E-08 | up | 3.93 | SMAD4 |
| 203077_s_at | 2.05E-09 | up | 2.50 | SMAD2 |
| 203140_at | 2.67E-08 | up | 4.67 | BCL6 |
| 203275_at | 3.53E-10 | up | 2.53 | IRF2 |
| 203749_s_at | 1.45E-09 | up | 2.78 | RARA |
| 204132_s_at | 7.42E-10 | up | 9.79 | FOXO3/B |
| 204790_at | 3.32E-06 | up | 2.69 | SMAD7 |
| 212550_at | 7.95E-10 | up | 2.70 | STAT5B |
| 214447_at | 2.18E-06 | down | -2.71 | ETS1 |
| 223287_s_at | 8.80E-09 | down | -4.59 | FOXP1 |
| 224889_at | 1.75E-10 | up | 8.90 | FOXO3 |
| 225223_at | 1.74E-06 | up | 2.84 | SMAD5 |
| 228188_at | 5.50E-07 | up | 4.34 | FOSL2 |
| 204959_at | 1.12E-06 | up | 5.58 | MNDA |
| 210029_at | 1.19E-05 | up | 3.98 | IDO1 |
| 225673_at | 1.62E-06 | up | 2.66 | MYADM |
| 202426_s_at | 5.52E-11 | up | 7.44 | RXRA |
| 244523_at | 7.85E-07 | up | 2.83 | MMD |
| 1565358_at | 1.29E-06 | down | 6.46 | RARA |

### TH17 related chemokine up-regulation during unstable angina

In this microarray analysis, we can see many chemokine genes are up-regulated during unstable angina. (Table 4) Most important of all, majority of these chemokine genes are TH17 related chemokines. The up-regulated chemokine as well as chemokine receptors include CXCL1, CCR1, CXCL6, CCR6, CCR3, DARC, CXCL16, CX3CR1, CXCL5, CKLF, IL8RB, IL8RA, IL8, S100A11, S100B, S100A9, S100P, LTB4R, and S100A12. Among these genes, CXCL1 is 8 fold up-regulated, CCR3 is 18 fold up-regulated, CXCL5 is 8 fold up-regulated, IL8RB is 38 fold up-regulated, and S100P is 50 fold up-regulated. Leukotriene B4 are mainly neutrophil chemoattractant. TH17 related chemokine or chemokine receptors include CXCL1, CXCL6, CXCL5, IL8RB, IL8RA, IL8, and S100 proteins.

**Table 4.** Chemokine

| Probe Set ID | Pvalue | Arrow | Fold | Gene |
| --- | --- | --- | --- | --- |
| 204470_at | 4.05E-11 | up | 7.82 | CXCL1 |
| 205098_at | 1.50E-05 | up | 4.00 | CCR1 |
| 206336_at | 4.69E-10 | up | 3.79 | CXCL6 |
| 208304_at | 4.74E-11 | up | 17.70 | CCR3 |
| 208335_s_at | 2.21E-06 | up | 3.16 | DARC |
| 215101_s_at | 2.78E-06 | up | 8.04 | CXCL5 |
| 219161_s_at | 4.00E-11 | up | 5.63 | CKLF |
| 223454_at | 4.20E-06 | up | 3.07 | CXCL16 |
| 1568934_at | 2.01E-06 | up | 2.65 | CX3CR1 |
| 207008_at | 1.99E-12 | up | 38.27 | IL8RB |
| 207094_at | 1.25E-14 | up | 12.51 | IL8RA |
| 211506_s_at | 3.18E-04 | up | 2.97 | IL8 |
| 202859_x_at | 0.02858 | up | 2.70 | IL8 |
| 208540_x_at | 2.08E-06 | up | 2.71 | S100A11/P |
| 209686_at | 0.022845 | up | 2.56 | S100B |
| 238909_at | 5.34E-06 | down | -2.87 | S100A10 |
| 203535_at | 4.97E-06 | up | 3.95 | S100A9 |
| 204351_at | 5.64E-10 | up | 50.29 | S100P |
| 205863_at | 1.63E-04 | up | 4.38 | S100A12 |
| 236172_at | 1.25E-09 | up | 3.39 | LTB4R |

### TH17 related cytokine up-regulation in unstable angina

During unstable angina, many TH17 related cytokines are up-regulated. (Table 5) Up-regulated TH17 related cytokines include IL-8 and IL-1B. It is worth noting that IL1B is 11 fold up-regulated. Down-regulated cytokines in unstable angina include IL-23A, IL-32, and TGFB1. It is important to know that IL1 beta is the key cytokine in TH17 anti-extracellular bacteria immunity. TGFB1 is down-regulated, suggesting that Treg cells are not important in unstable angina. Significant up-regulation of IL-1 receptors (10 fold increase for IL1R1 and 37 fold increase for IL1R2) suggests that TH17 immunological pathway is activated. Another TH17 player, IL6R is also up-regulated in the analysis. IFNGR1 is 10X down-regulated, suggesting traditional TH1 immunity is not activated in unstable angina.

**Table 5.** Cytokine&receptors

| Probe Set ID | Pvalue | Arrow | Fold | Gene |
| --- | --- | --- | --- | --- |
| 1568513_x_at | 5.20E-07 | down | -3.15 | IL23A |
| 202859_x_at | 0.02858 | up | 2.70 | IL8 |
| 203828_s_at | 2.70E-07 | down | -4.22 | IL32 |
| 205067_at | 2.91E-09 | up | 11.20 | IL1B |
| 203085_s_at | 2.46E-12 | down | -4.48 | TGFB1 |
| 205016_at | 3.08E-13 | up | 12.56 | TGFA |
| 202948_at | 3.01E-13 | up | 10.60 | IL1R1 |
| 204116_at | 1.72E-13 | down | -4.50 | IL2RG |
| 205403_at | 1.14E-14 | up | 37.11 | IL1R2 |
| 206618_at | 3.97E-05 | up | 2.90 | IL18R1 |
| 209575_at | 6.50E-10 | up | 3.09 | IL10RB |
| 210904_s_at | 6.93E-13 | up | 10.39 | IL13RA1 |
| 226333_at | 4.08E-13 | up | 2.73 | IL6R |
| 225661_at | 1.28E-12 | up | 5.96 | IFNAR1 |
| 242903_at | 1.66E-09 | down | -10.03 | IFNGR1 |

### Anti-bacterial complement up-regulation during unstable angina

Complements are important for anti-bacterial innate TH17 immunity. Here, we show that majority of complement genes are up-regulated during unstable angina. (Table 6) These complement genes include CD59, CD55, CFD, C4BPA, C4A/B, CD46, C1QBP, ITGAX, C1RL, C5AR1, and CR1/1L. It means that the whole complement machinery is up-regulated during unstable angina.

**Table 6.**
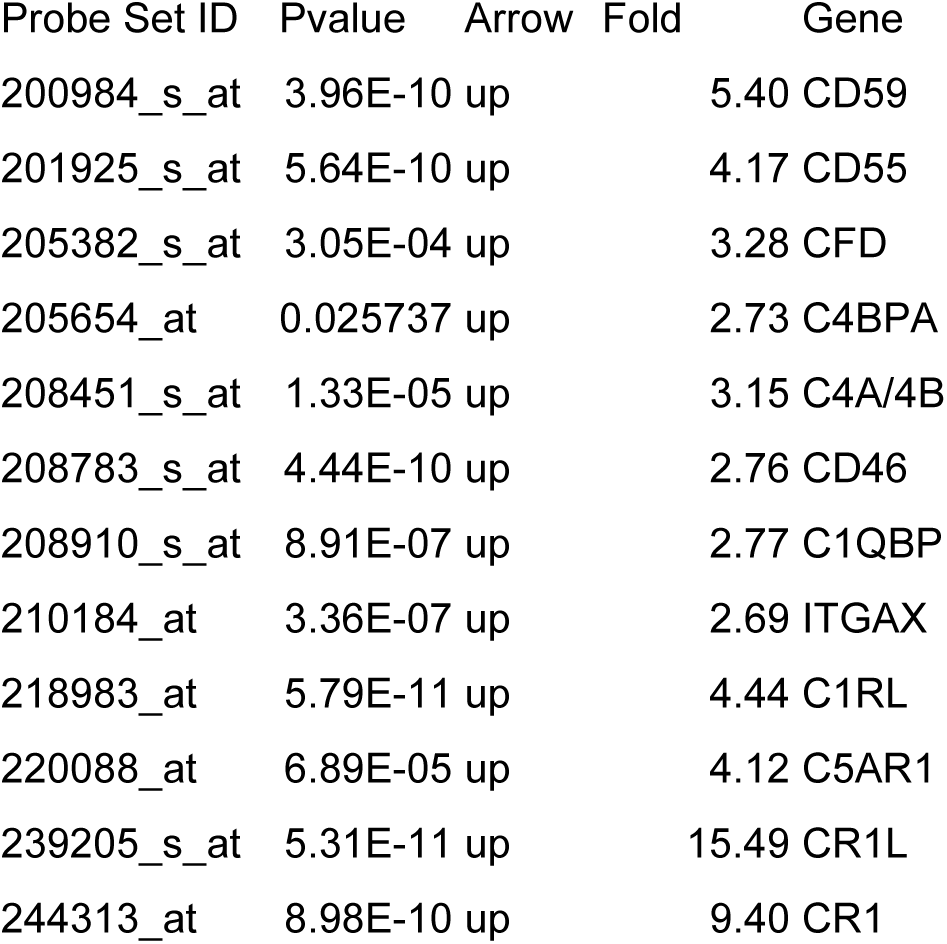
Complement

### Other anti-bacterial related gene up-regulation in unstable angina

In table 7, many other important anti-bacterial genes are up-regulated during unstable angina. These genes include CSF3R, FPR1, CSF2RB, SCARF1, CSF2RA, DEFA4, NCF4, NCF2, FPR2, MRC2, and PTX3. It is worth noting that CSF2RB is 10 fold up-regulated, DEFA4 is 11 fold up-regulated, and FPR2 is 38 fold up-regulated. These neutrophil or anti-bacterial related genes suggest the activation of TH17 host immunological pathways in unstable angina.

**Table 7.** anti-bacterial genes

| Probe Set ID | Pvalue | Arrow | Fold | Gene |
| --- | --- | --- | --- | --- |
| 203591_s_at | 5.40E-05 | up | 2.95 | CSF3R |
| 205119_s_at | 1.17E-07 | up | 8.65 | FPR1 |
| 205159_at | 4.60E-08 | up | 10.71 | CSF2RB |
| 206995_x_at | 9.73E-09 | up | 5.09 | SCARF1 |
| 207085_x_at | 1.37E-06 | up | 3.86 | CSF2RA |
| 207269_at | 9.38E-06 | up | 11.52 | DEFA4 |
| 207677_s_at | 4.04E-08 | up | 5.47 | NCF4 |
| 209949_at | 2.90E-05 | up | 2.76 | NCF2 |
| 210772_at | 3.39E-13 | up | 38.75 | FPR2 |
| 37408_at | 1.83E-05 | up | 2.61 | MRC2 |
| 206157_at | 9.63E-08 | up | 3.04 | PTX3 |

### Glycolytic genes up-regulation during unstable angina

In table 8, the majority of glycolytic pathway is up-regulated in unstable angina. Hypoxia can drive the activation of the anaerobic glycolysis. Thus, it is reasonable that glycolytic enzymes are up-regulated during the hypoxia status of the attack of unstable angina. These up-regulated genes include PGK1, PFKFB3, PYGL, BPGM, PDK3, PFKFB2, ENO1, and PGK1. It is worth noting that BPGM(2,3-bisphosphoglycerate mutase) is 19 fold up-regulated. The product of BPGM is 2,3-bisphosphoglycerate, which combines with hemoglobin, can cause a decrease in affinity for oxygen. Thus, the presence of 2,3-bisphosphoglycerate helps oxyhemoglobin to unload oxygen to help unstable angina patients.

**Table 8.** Glycolytic enzymes

| Probe Set ID | Pvalue | Arrow | Fold | Gene |
| --- | --- | --- | --- | --- |
| 217356_s_at | 1.45E-07 | up | 3.49 | PGK1 |
| 202464_s_at | 1.88E-05 | up | 3.85 | PFKFB3 |
| 202990_at | 4.01E-07 | up | 4.85 | PYGL |
| 203502_at | 2.42E-08 | up | 19.11 | BPGM |
| 206348_s_at | 4.04E-08 | up | 2.49 | PDK3 |
| 209992_at | 1.30E-07 | up | 2.55 | PFKFB2 |
| 217294_s_at | 9.10E-05 | up | 3.21 | ENO1 |
| 217356_s_at | 1.45E-07 | up | 3.49 | PGK1 |

### H^+^-ATPase gene up-regulation during unstable angina

In table 9, we can see many H^+^-ATPases are up-regulated during unstable angina. These genes include ATP6V0C, ATP6V1B2, ATP6V0E1, ATP6V1C1, ATP6V0A2, and ATP6V1D. These ATPases are proton pumps which can generate H^+^ by using ATP. In addition, several carbonic anhydrases are also up-regulated including CA1 and CA4. Carbonic anhydrases can catalyze H_2_CO_3_ formation from CO_2_ and H_2_O. Thus, acidosis can result during the attack of unstable angina.

**Table 9.** ATPase

| Probe Set ID | Pvalue | Arrow | Fold | Gene |
| --- | --- | --- | --- | --- |
| 200954_at | 7.36E-08 | up | 3.46 | ATP6V0C |
| 201089_at | 2.00E-05 | up | 3.34 | ATP6V1B2 |
| 202872_at | 1.07E-10 | up | 6.46 | ATP6V1C1 |
| 208899_x_at | 3.59E-08 | up | 2.72 | ATP6V1D |
| 214149_s_at | 8.54E-11 | up | 7.66 | ATP6V0E1 |
| 205950_s_at | 5.00E-08 | up | 42.28 | CA1 |
| 206209_s_at | 1.82E-12 | up | 8.94 | CA4 |

### Coagulation gene up-regulation during unstable angina

In table 10, majority of coagulation related genes are up-regulated in peripheral leukocytes of unstable angina patients. These genes include THBS1, THBD, F2R, F5, F8, F2RL1, PROS1, GP5, PTAFR, PLAUR, TFP1, ITGB3, HPSE, GP6, PEAR1, and TBXAS1. Most important of all, F2RL1 is 18 fold up-regulated.

**Table 10.** Coagulation genes

| Probe Set ID | Pvalue | Arrow | Fold | Gene |
| --- | --- | --- | --- | --- |
| 201110_s_at | 5.89E-08 | up | 6.73 | THBS1 |
| 203887_s_at | 2.66E-10 | up | 13.16 | THBD |
| 203989_x_at | 3.03E-05 | up | 3.24 | F2R |
| 205756_s_at | 1.71E-07 | up | 2.89 | F8 |
| 207808_s_at | 6.13E-06 | up | 3.83 | PROS1 |
| 207815_at | 9.67E-06 | up | 8.68 | PF4V1 |
| 207926_at | 1.53E-05 | up | 2.54 | GP5 |
| 211661_x_at | 1.44E-06 | up | 5.10 | PTAFR |
| 211924_s_at | 1.35E-07 | up | 4.40 | PLAUR |
| 213258_at | 9.09E-06 | up | 3.44 | TFPI |
| 213506_at | 1.11E-10 | up | 18.07 | F2RL1 |
| 215240_at | 1.86E-07 | up | 4.26 | ITGB3 |
| 220336_s_at | 3.37E-04 | up | 2.51 | GP6 |
| 222881_at | 1.96E-06 | up | 2.97 | HPSE |
| 228618_at | 2.78E-05 | up | 2.61 | PEAR1 |
| 231029_at | 2.14E-09 | up | 4.81 | F5 |
| 236345_at | 1.63E-13 | up | 8.19 | TBXAS1 |

### RBC related genes are up-regulated during unstable angina

In table 11, majority of RBC related genes are up-regulated during unstable angina. These up-regulated genes include EPB41, GYPE, ANK1, NFE2, ALAS2, GYPA, HBG1/HBG2, EPOR, RHCE/RHD, GYPB, HEBP1, ERAF, HBQ1, HEMGN, HBM, and HBD. Borderline down-regulated gene includes ANKRD10. It is worth noting that many hemoglobin genes are up-regulated including HBG1, HBG2, HBQ1, HBM, and HBD. Among them, HBG1/2 is 20 fold up-regulated, HBM is 60 fold up-regulated, HEMGN is 49 fold up-regulated, HBQ1 is 18 fold up-regulated, ERAF is 33 fold up-regulated, GYPA is 27 fold up-regulated, and ALAS2 is 36 fold up-regulated.

**Table 11.**
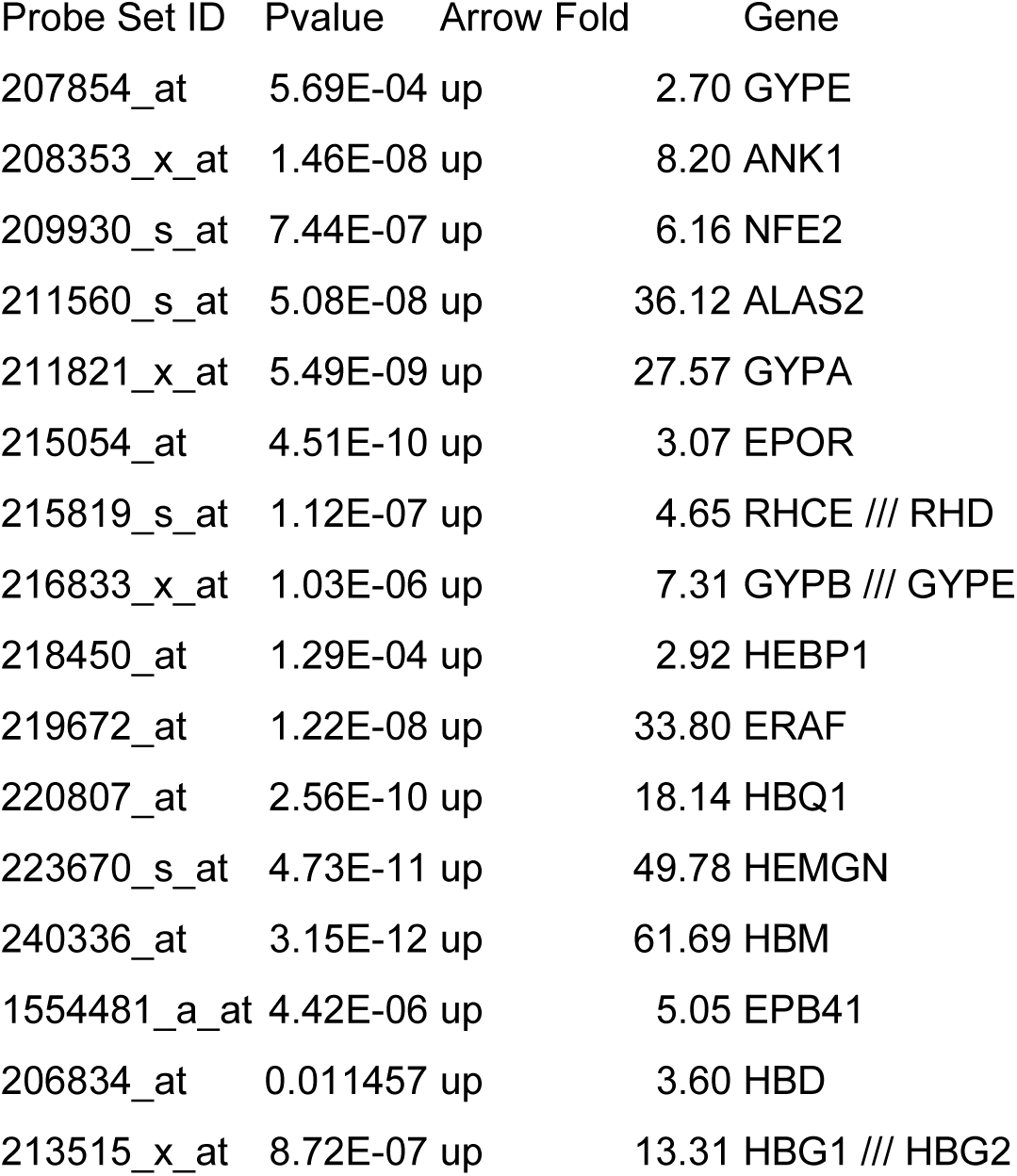
RBC genes

### IL-17 centered top regulator effect network in pathway analysis

By using Ingenuity Pathway Analysis, IL17-STAT3 immunological pathway is the number one over-represented regulator pathway in the whole blood RNA samples in unstable angina patients compared to control. (Figure 3) Thus, Th17 immunological reaction plays a central role in the pathophysiology of unstable angina. Unstable angina is closely related to Th17 immunity. Besides, the top toxicology function of unstable angina is represented by myocardial infarction which proves that this study’s validity. (Figure 4) The activated top upstream regulator is TNF network including IL1B, STAT1, STAT3, NFkB, JUN, and RELA which are also matching Th17 immunological pathway. (Figure 5) The top network is RXRA centered pathway. RXRA is one of the key transcription factor in Th17 immunity. (Figure 6) The top five functions of unstable angina are: cell death and survival, infectious disease, immunological disease, inflammatory disease, organismal injury and abnormalities. (Figure 7) Thus, immune reaction plays a key role in the pathogenesis of unstable angina. Finally, the top activated molecular pathways of unstable angina include B cell receptor signaling, IL8 signaling, and NFAT immune regulation signaling. (Figure 8-11). IL8 is the key chemokine in Th17 immunity. These also suggest that immunity as well as Th17 immunological pathway plays an important role in unstable angina.

**Figure 3.**
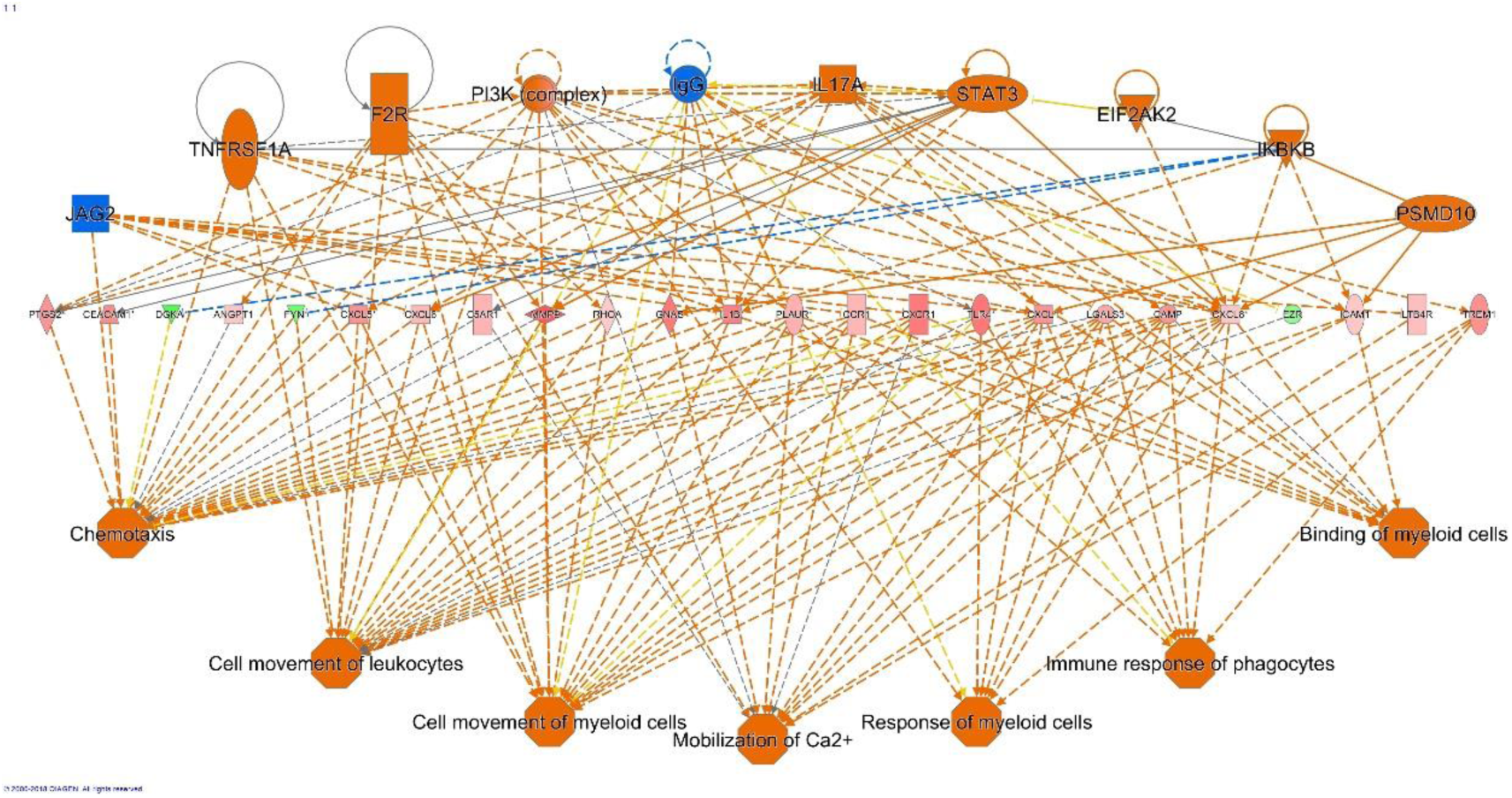
Top effector regulator pathway in unstable angina. The first row includes the top effector regulator. The second role includes the significantly expressed genes in our dataset. The third law is the effector function. Red color means up-regulation and green color means down-regulation.

**Figure 4.**
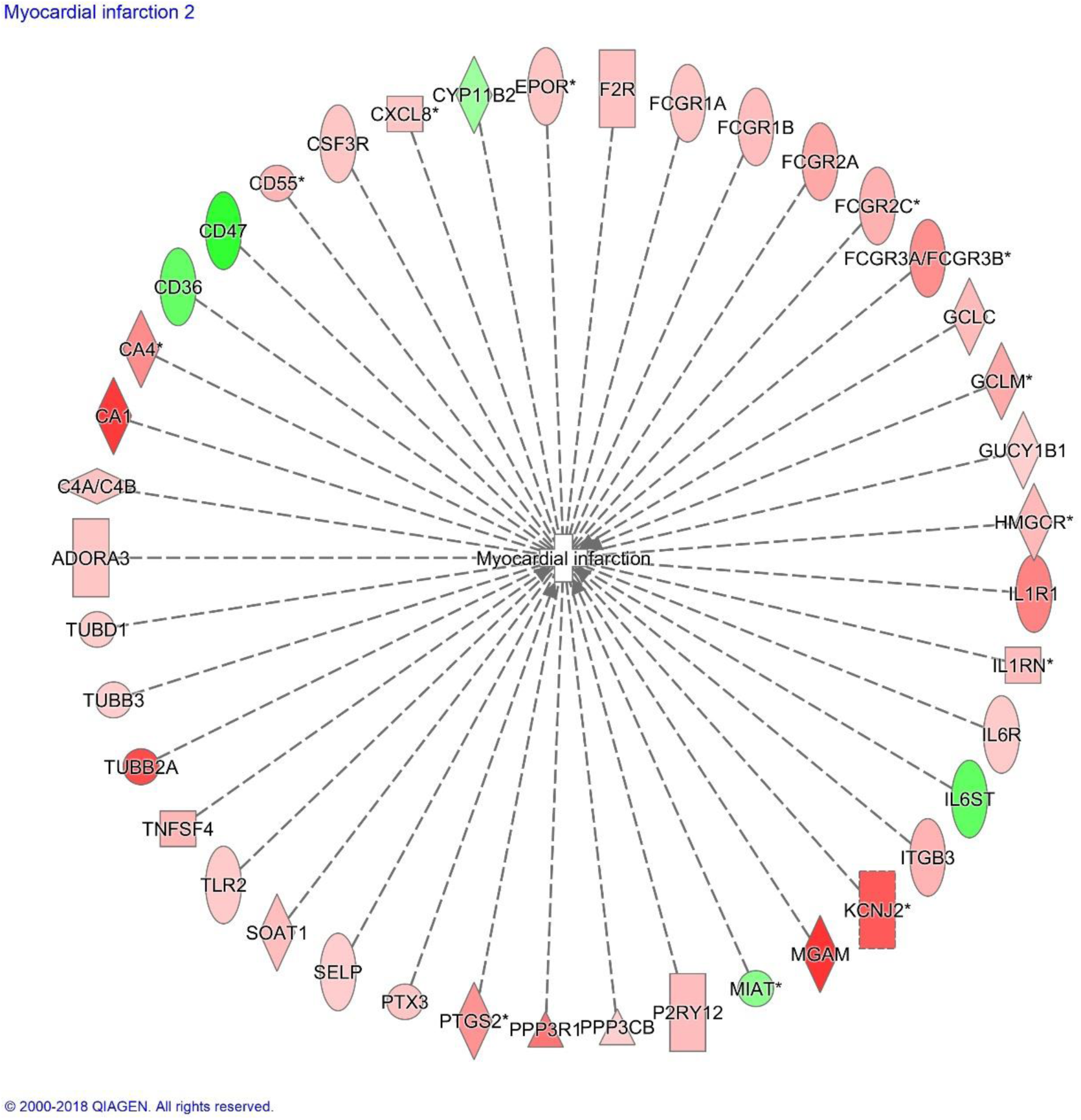
Top toxicological function in unstable angina. The central node is the predicted top toxicological function. The peripheral genes are the significantly expressed genes related to the central toxicological function. Red color means up-regulation and green color means down-regulation.

**Figure 5.**
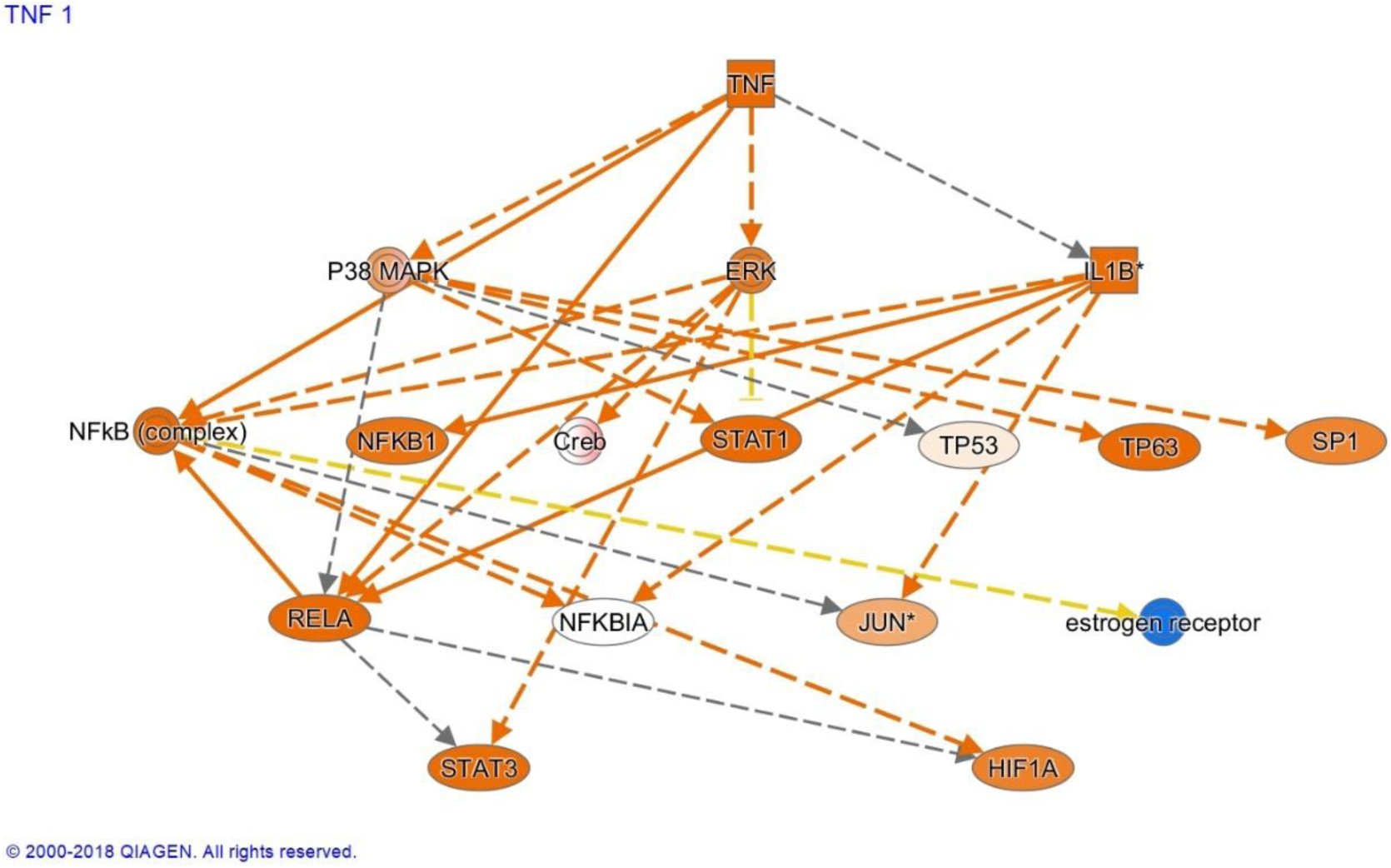
Top upstream mediator in unstable angina. TNF is the top upstream mediator in our dataset. Stronger red color means more highly expressed, and lesser red color means less highly expressed. All other genes are downstream of TNF.

**Figure 6.**
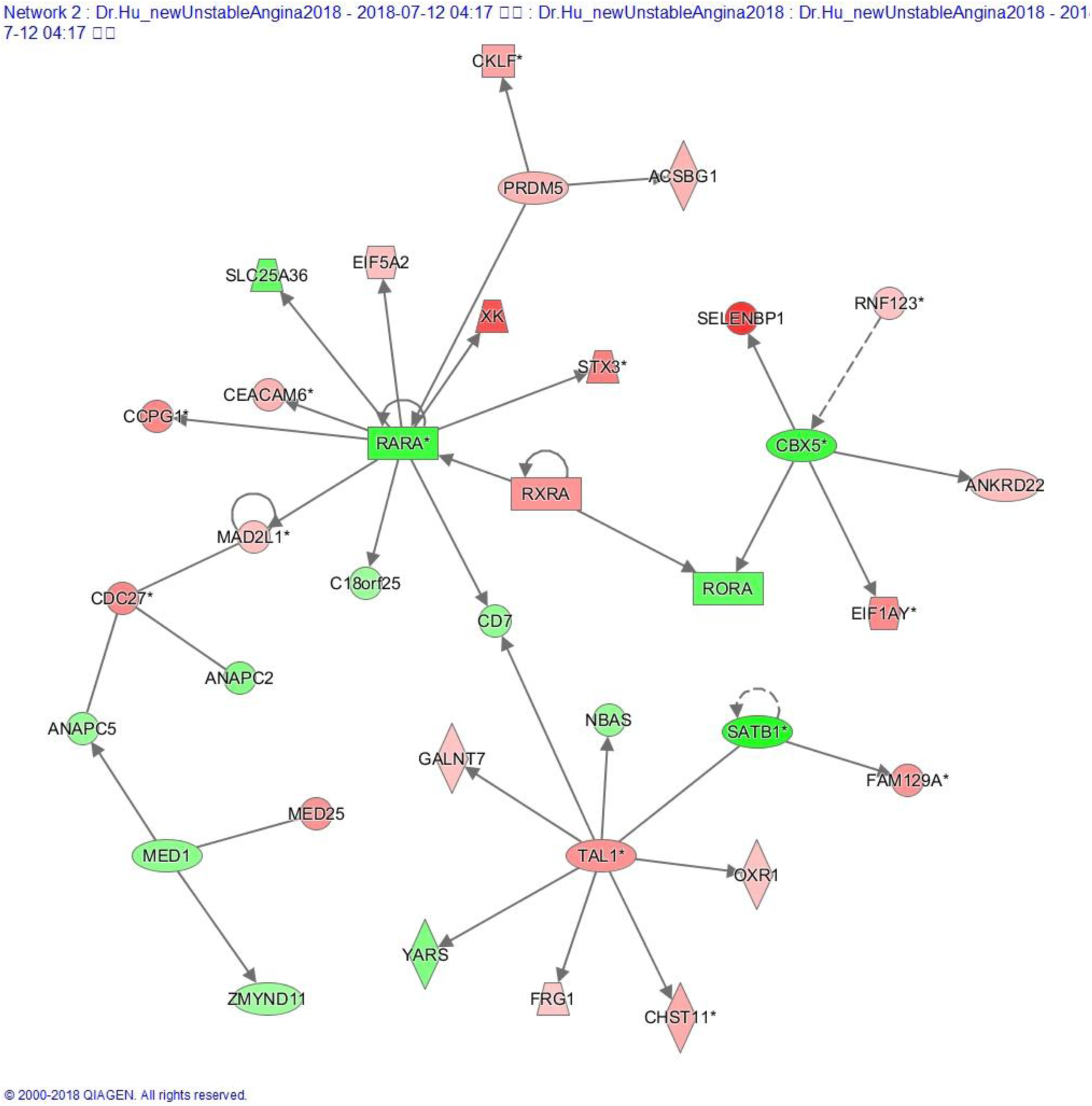
Top represented network in unstable angina. The most represented network in unstable angina dataset is the RXRA-RARA-RORA centered network. RXRA is up-regulated in our analysis. RARA has two probe-sets in our data; one is up-regulated, and the other one is down-regulated. IPA selected down-regulated one for this graph due to higher fold change. However, the up-regulated one has more significant P value.

**Figure 7.**
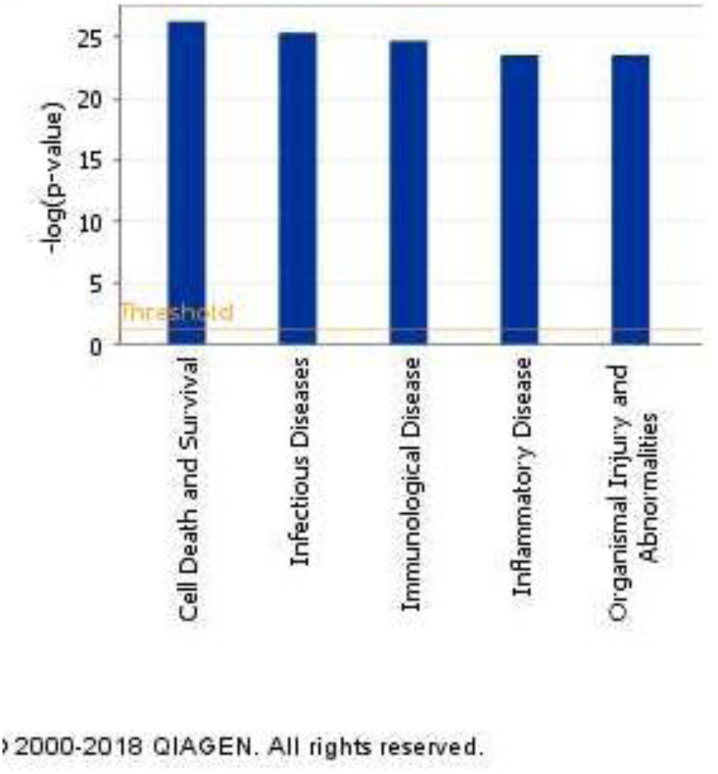
Top five functions in unstable angina. The most significantly expressed physiological functions are displayed by the rank of -log(P value).

**Figure 8.**
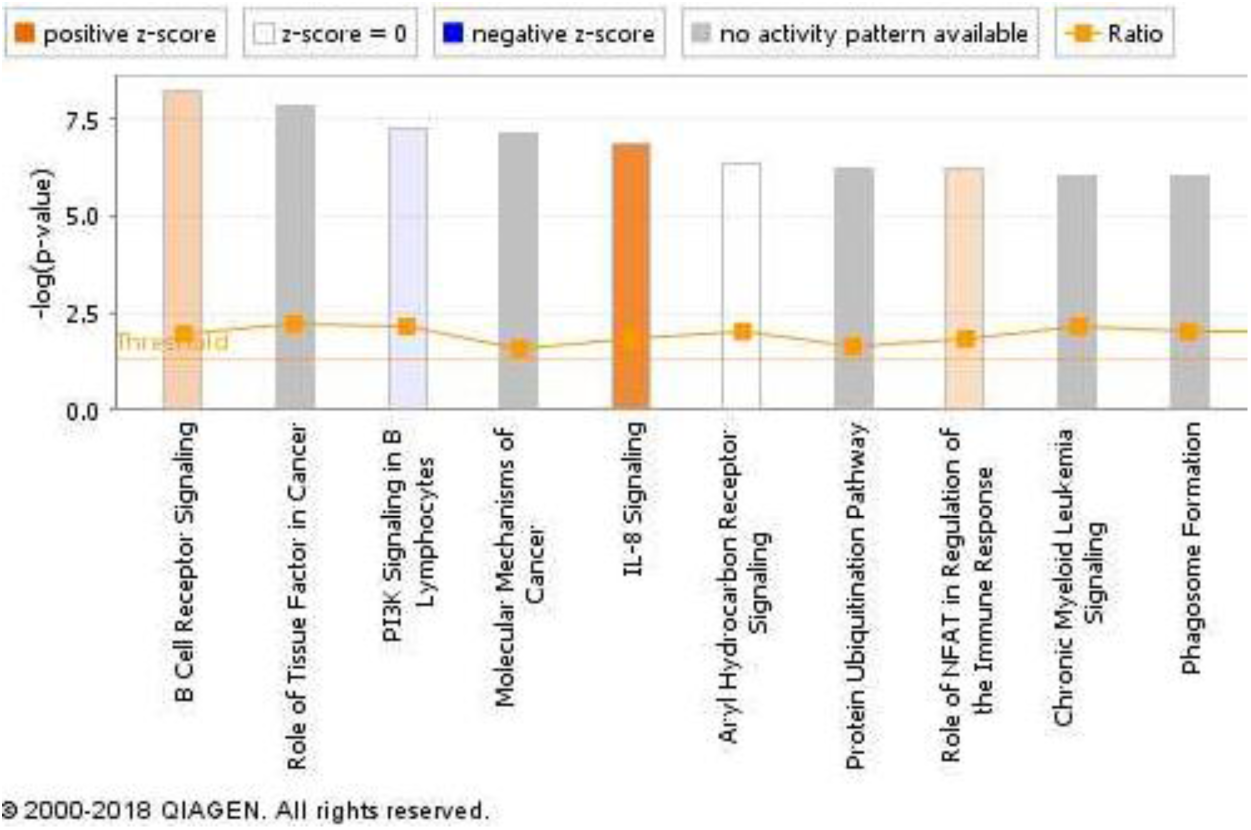
Top molecular pathways in unstable angina. The most significantly expressed molecular pathways are displayed by the rank of -log(P value). Positive z-score (red color) means the molecular pathway is activated, negative z-score (blue color) means the molecular pathway is inhibited, and the zero z-score or grey color means no available activity pattern.

**Figure 9.**
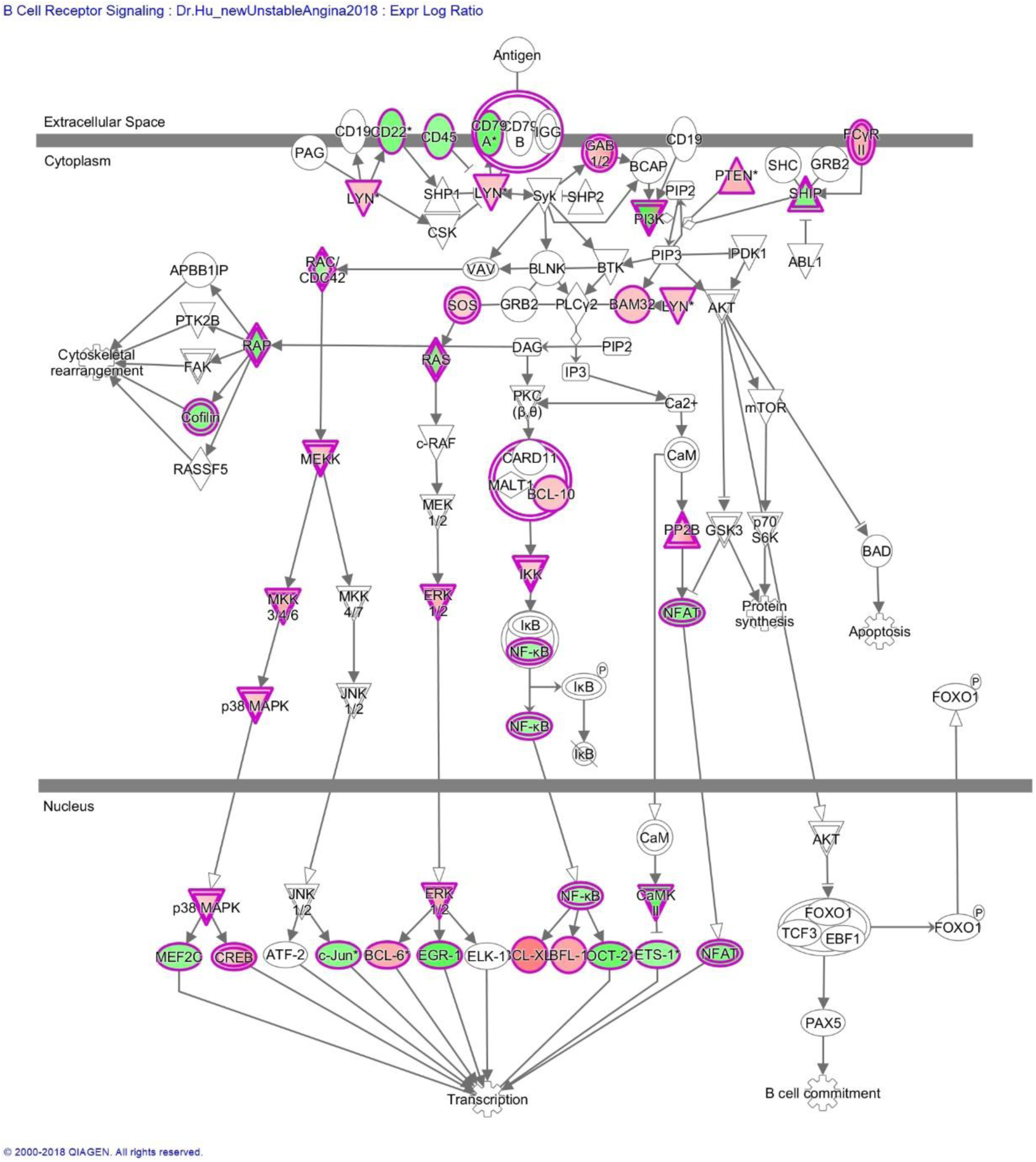
B cell receptor signaling activation in unstable angina. Red color means up-regulation and green color means down-regulation.

**Figure 10.**
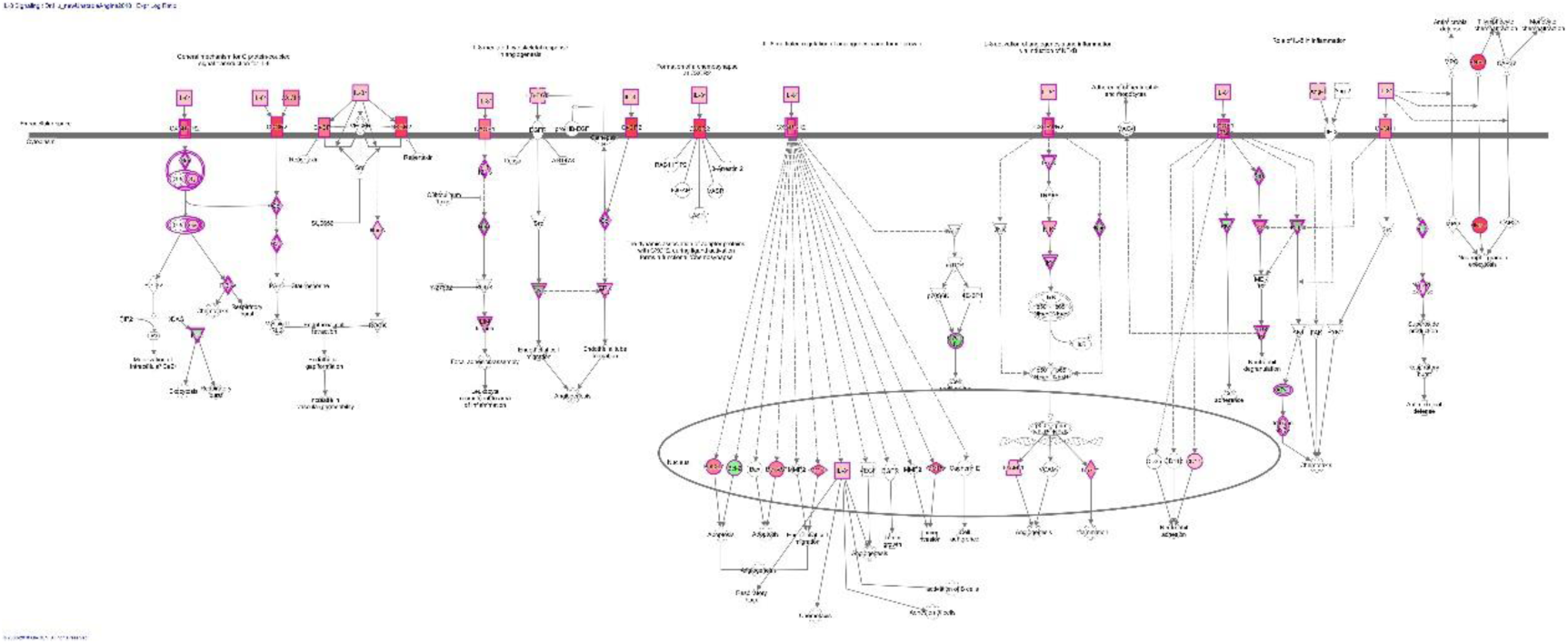
IL8 signaling activation in unstable angina. Red color means up-regulation and green color means down-regulation.

**Figure 11.**
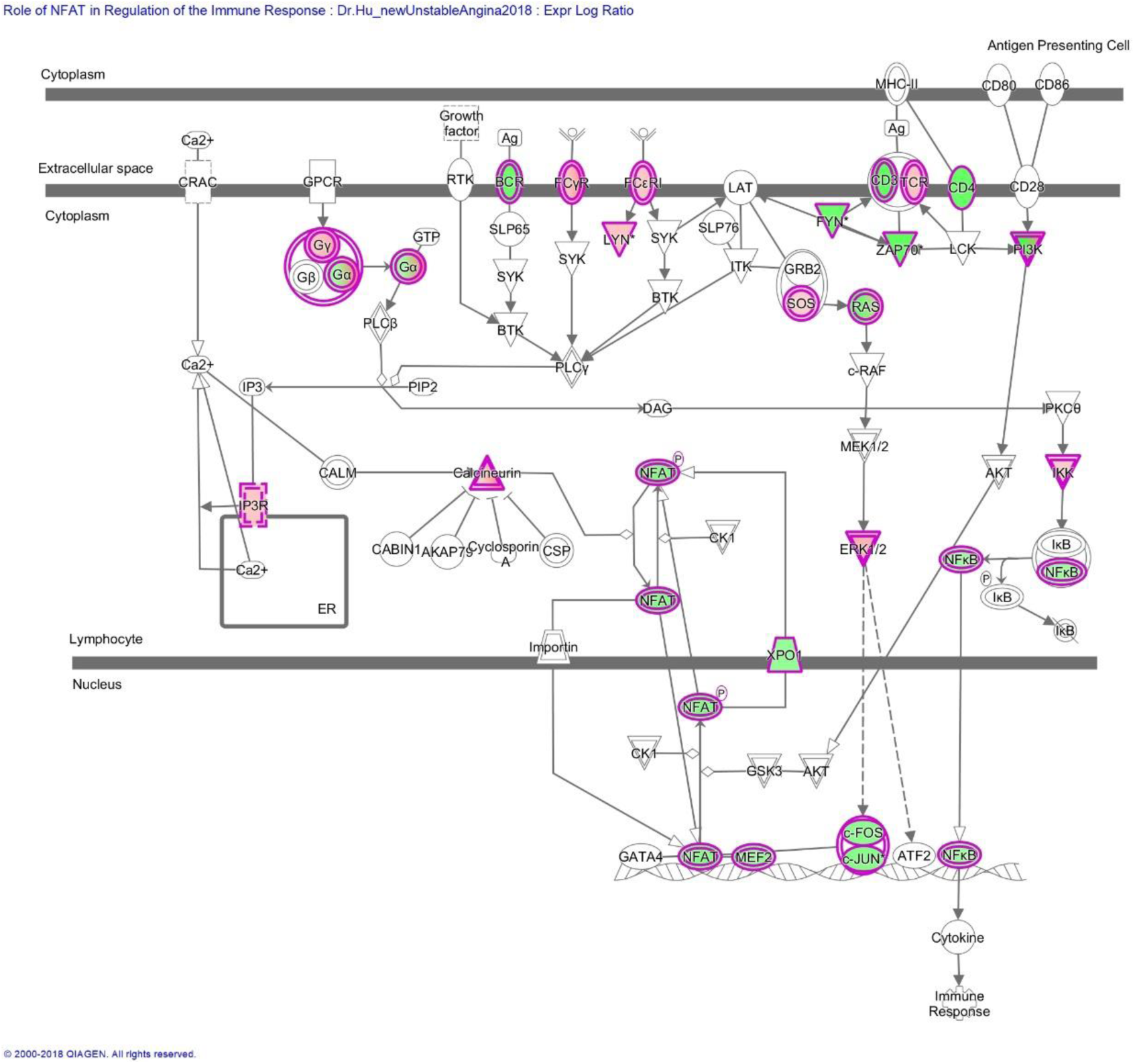
NFAT mediator signaling activation in unstable angina. Red color means up-regulation and green color means down-regulation.

## Discussion

Atherosclerosis is a very common and detrimental disease, especially in many developed countries. It is usually among the top three leading causes of mortality. Inflammatory process is thought to be related to the etiology of atherosclerosis. C reactive protein (CRP) is found to be related to the course of atherosclerosis. Abnormal T cell activation has been found in atherosclerosis.(Liuzzo, Kopecky et al. 1999) However, detailed relationship between inflammation and atherosclerosis is remained to be unlocked. Unstable angina is a clinical manifested syndrome of atherosclerosis. Thus, I use microarray analysis of peripheral leukocytes to study the relationship of specific inflammation and unstable angina.

Host immunity against extracellular bacteria induces TH17 immune response. Helicobacter pylori infection is also related to atherosclerosis. Thus, anti-extracellular TH17 immunity is also likely to be activated during atherosclerosis(Ng, Burris et al. 2011). In addition, TH17 immunological pathways are also related to important risk factors for atherosclerosis: Diabetes mellitus and hypertension.(Gao, Jiang et al. 2010, Xie, Wang et al. 2010) In addition, CRP, the TH17 inflammatory factor, is elevated in atherosclerosis. And, deficiency of Treg cells promotes the progression of atherosclerosis.(Gotsman, Grabie et al. 2006, Mor, Planer et al. 2007) Besides, TH2 and THαβ immunological pathway activation such as IL-10 up-regulation can reduce the development of atherosclerosis.(Liuzzo, Kopecky et al. 1999, Pinderski 2002) These evidences all suggest that TH17 immunity plays an important role in the pathophysiology of atherosclerosis.

In this study, I find out the evidences that immunity plays a very important role in the pathogenesis of unstable angina. First of all, anti-bacterial Toll-like receptors are up-regulated during unstable angina. These Toll-like receptors include TLR 1,2,4,8. It is worth noting that TLR 1,2,4 are for triggering anti-extracellular bacteria TH17 immunity. Thus, TH17 immunity is initiated during unstable angina.

In addition, heat shock proteins which are responsible for triggering anti-bacterial immunity are also up-regulated during unstable angina attack. Heat shock protein 70 (HSP70) can bind to Toll-like receptor 2 and Toll-like receptor 4 to trigger TH17 immunity. In this study, I find out that HSP70 genes are up-regulated including HSPA1A and HSPA6. This means anti-bacterial immunity is activated during unstable angina.

Then, we look at immune-related transcription factor expression in unstable angina. Strikingly, we find out many TH17 related immune transcription factor up-regulation in unstable angina patients’ peripheral leukocytes. STAT1, STAT5B, and BCL6 are the key downstream transcription factors of follicular helper T cells. Follicular helper T cells activity mean the adaptive immunity is initiated. In addition, SMAD proteins are major effector genes of TGF beta signaling. TGF beta is the key cytokine to initiate TH17 immunity. Thus, TH17 is likely to be activated during unstable angina. In addition, TH17 negative regulator, ETS1, is down-regulated in unstable angina(Moisan, Grenningloh et al. 2007). Retinoic acid related transcription factors, which promote TH1 and TH17 immunity, are up-regulated during unstable angina.(Mucida, Park et al. 2007)

Chemokine genes are differentially regulated in unstable angina. Most important of all, TH17 related chemokine and chemokine receptor genes are up-regulated. These strikingly up-regulated chemokine related genes include CXCL1, CCR3, CXCL5, IL8RB, IL8RA, and S100P. CXCL1, CXCL5, and S100P are TH17 immunity related chemokines. And, IL8RB and IL8RA are TH17 related chemokine receptor. Besides, leukotriene B4 related gene is also up-regulated during unstable angina. This also shows that TH17 anti-bacterial immunity is activated at unstable angina.

Then, we look at cytokine and cytokine receptor gene expression profiles of peripheral leukocytes during unstable angina. We find out that TGFA, IL1B, and IL8 are up-regulated. Among them, IL8 and IL1B are key TH17 related cytokines. Besides, many cytokine receptors are also differentially regulated at unstable angina. These up-regulated cytokine receptors include IL-1R, IL6R, and TGFB1R. TGFB and IL-6 are driven cytokines for TH17 immunity. IL8 is the main neutrophil chemotaxis mediator. IL1B is a key effector cytokine in TH17 immunity to activate neutrophils.

Complements are important in mediating host killing of extracellular bacteria. In this study, we can see that majority of complement machinery is activated including CD59, CD55, CFD, C4BPA, CR1, C4A/B, CD46, C1QBP, ITGAX, C1RL, and C5AR1. This means anti-bacterial TH17 immunity is triggered during unstable angina. Besides, other important anti-bacterial host defense genes are also up-regulated. These up-regulated anti-bacterial genes include CSF3R, FPR1, CSF2RB, SCARF1, CSF2RA, DEFA4, NCF4, NCF2, FPR2, MRC2, and PTX3. These genes include neutrophil and macrophage growth factors, pattern recognition receptor, defensin, neutrophil cytosolic factors, and pentraxin. These all suggest that TH17 anti-bacterial immunity is activated during unstable angina.

By using microarray analysis, we can also understand the complications of unstable angina. Glycolysis, acidosis, hypoxia, and coagulation anomaly are frequently reported in unstable angina. First of all, we look at the glycolytic mediating enzymes in unstable angina. Strikingly, we find out all glycolytic enzyme genes are up-regulated including PGK1, PFKFB3, PYGL, BPGM, PDK3, PFKFB2, ENO1, and PGK1. Among them, BPGM is 19 fold up-regulated. BPGM can help to unload oxygen from oxy-hemoglobins. Thus, it can help to alleviate the hypoxic syndrome during unstable angina. It can be explained as a host compensatory mechanism responding to the attack of unstable angina. More, we find out that many H^+^ATPase genes are up-regulated during unstable angina including ATP6V0C, ATP6V1B2, ATP6V0E1, ATP6V1C1,and ATP6V1D. Several carbonic anhydrase enzyme genes are also up-regulated including CA1 and CA4. In my previous malaria genomic research, I find out a co-expression of glycolytic enzymes and ATPases. Here, I also find out acidosis at unstable angina can be due to up-regulation of these proton pumps (H^+^-ATPase) and H_2_CO_3_ producing carbonic anhydrases.

Finally, we look at RBC and platelet related gene expression profiles during unstable angina. Strikingly, we find out that majority of RBC and platelet related genes are up-regulated during unstable angina. Up-regulated platelet/coagulation related genes include THBS1, THBD, F2R, F5, F8, F2RL1, PROS1, GP5, PTAFR, PLAUR, TFP1, ITGB3, HPSE, GP6, PEAR1, and TBXAS1. Platelet function is triggering blood coagulation, and overactivation of coagulation related genes can be related to the pathophysiology of atherosclerosis induced unstable angina with coronary artery occlusion. Up-regulated RBC related genes include EPB41, GYPE, ANK1, HP/HPP, NFE2, ALAS2, GYPA, HBG1/HBG2, EPOR, RHCE/RHD, HEBP1, HBQ1, HEMGN, HBM, and HBD. We can see many hemoglobin genes are up-regulated at unstable angina. Unstable angina is caused by lack of oxygen support for cardiac muscle. Thus, up-regulated RBC related genes including hemoglobin genes can help to alleviate the hypoxia status during unstable angina. It can be viewed as a host compensatory mechanism at the attack of unstable angina.

By using pathway analysis, we can see the top regulator network is IL17-STAT3 centered pathway. Thus, it confirms that TH17 immunological pathway plays a vital role in the pathophysiology of unstable angina. The upstream mediator is TNF, the top network is RXRA, and the top molecular pathway is IL8 signaling. Besides, cell death and survival, immune response represents the top functions in unstable angina. These also suggest that Th17 immunity is key mediator in the pathogenesis of unstable angina.

In summary, microarray analysis of peripheral leukocytes from unstable angina patients can help to explain the pathogenesis of unstable angina. From the above evidences, unstable angina can be viewed as a TH17 inflammatory disorder with up-regulated TH17 chemokines, cytokines, transcription factors, Toll-like receptors, heat shock proteins, complements, leukotrienes, prostaglandins, defensins, pentraxins, and other effector molecules. In addition, overactivation of platelet related genes, glycolytic enzymes, and H^+^-ATPases can help to explain the pathophysiology of unstable angina with coagulation hyperactivity and metabolic acidosis. Finally, up-regulation of RBC related genes should be a host compensatory mechanism to increase oxygen delivery during unstable angina.

## Author’s information

Wan-Chung Hu is a MD from College of Medicine of National Taiwan University and a PhD from vaccine science track of Department of International Health of Johns Hopkins University School of Public Health. He is a postdoctorate in Genomics Research Center of Academia Sinica, Taiwan. His previous work on immunology and functional genomic studies were published at *Infection and Immunity* 2006, 74(10):5561, *Viral Immunology* 2012, 25(4):277, and *Malaria Journal* 2013,12:392. He proposed THαβ immune response as the host immune response against viruses.

